## Supplemental Data 1 for "How human runners regulate footsteps on uneven terrain"

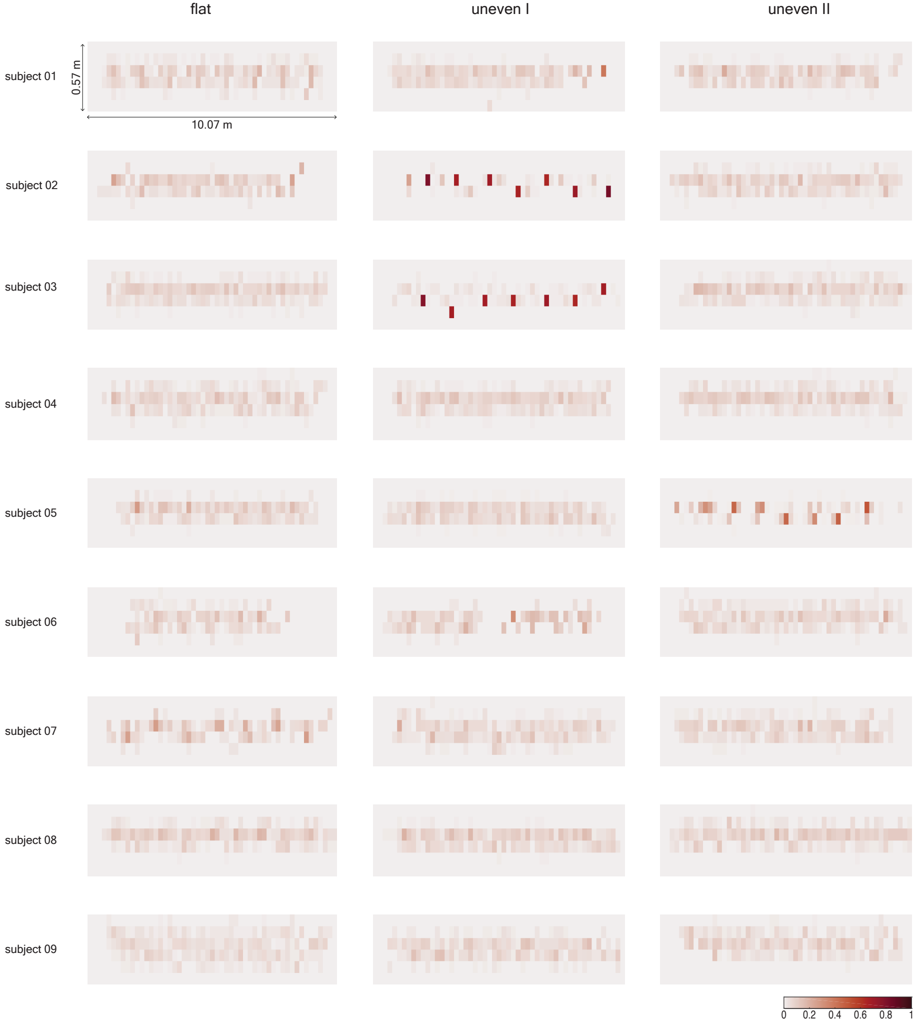


**Figure 3—figure supplement 1. Subject-wise foot placement patterns.** Heatmaps of the foot placement index for all subjects on each terrain. Each cell has an area of 190 mm × 95 mm, with the longer side of the rectangle along the length of the track. Colour bar at the bottom right of the figure shows the value of $p_{i,j}$.
