## Supplemental Data 2 for "How human runners regulate footsteps on uneven terrain"

**Figure 3—table supplement 1.** Footstep counts for each subject on all terrain.

| subject | flat | uneven I | uneven II |
|---|---|---|---|
| 1 | 363 | 321 | 448 |
| 2 | 228 | 78 | 473 |
| 3 | 473 | 131 | 436 |
| 4 | 373 | 471 | 519 |
| 5 | 366 | 557 | 109 |
| 6 | 224 | 160 | 327 |
| 7 | 218 | 398 | 442 |
| 8 | 503 | 489 | 479 |
| 9 | 477 | 390 | 392 |
