## Supplemental Data 4 for "How human runners regulate footsteps on uneven terrain"

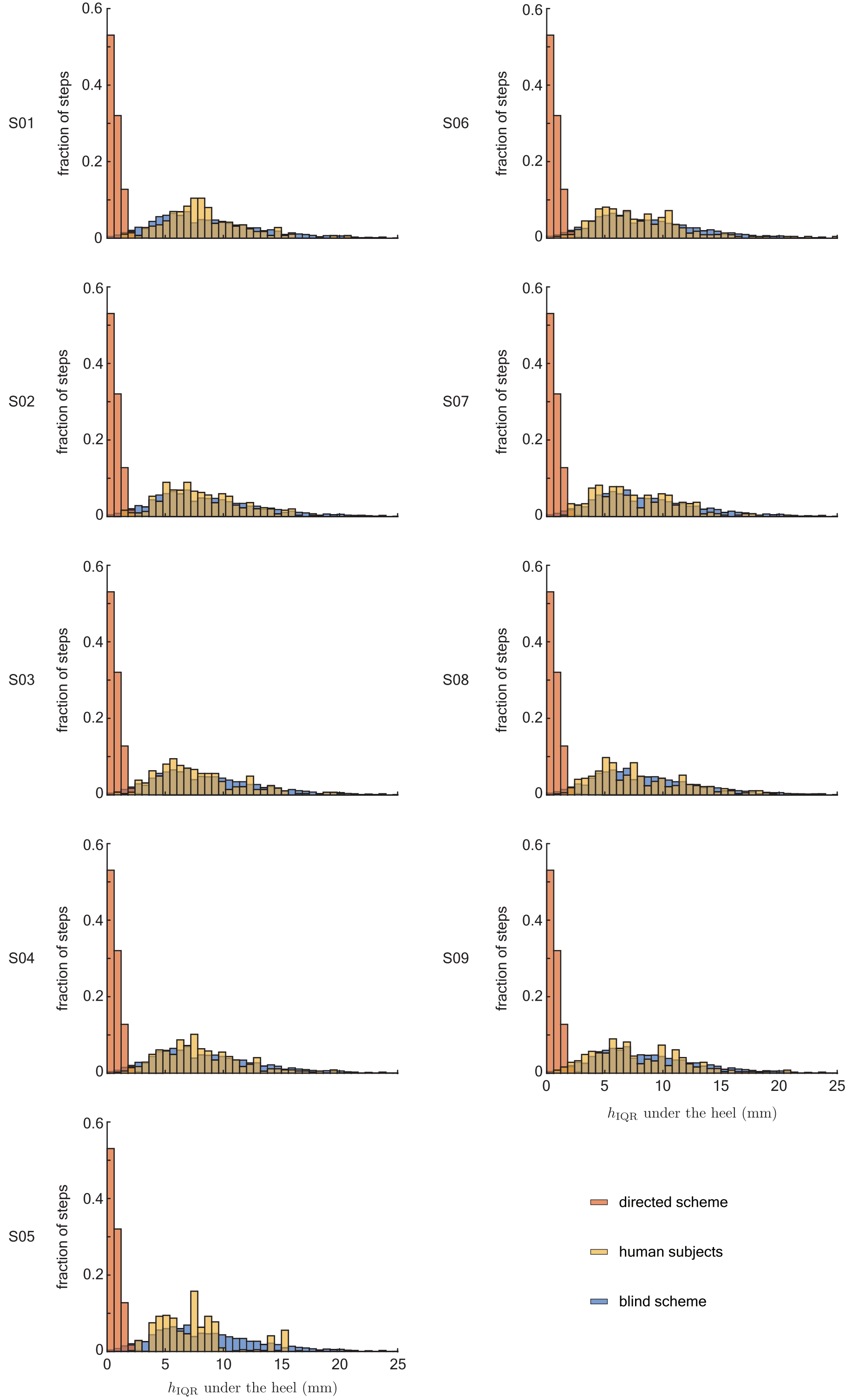


**Figure 5—figure supplement 2. Subject-wise foot placement analysis on uneven II.** Histograms of the interquartile range of heights (IQR) at the footstep locations for the directed sampling scheme (red), each subject from the experiments (yellow), and the blind sampling scheme (blue).
