## Supplemental Data 5 for "How human runners regulate footsteps on uneven terrain"

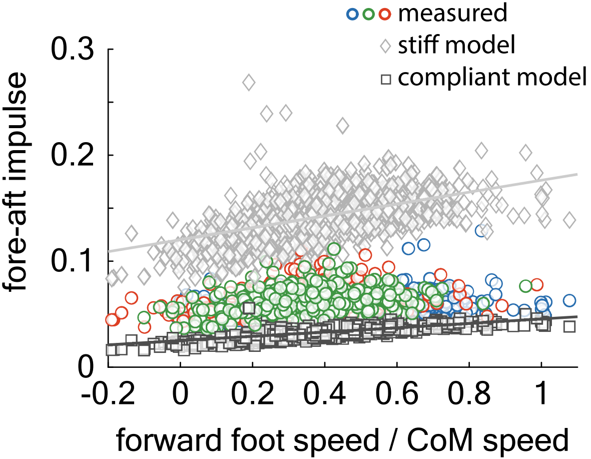


**Figure 6—figure supplement 1. Detailed results of the collision analysis.** Measured $J_y^*/(mv_y)$ (blue circles: flat, red circles: uneven I, green circles: uneven II) and calculated $J_y^*/(mv_y)$ values for the collision model with a compliant leg (dark grey squares) and rigid leg (light gray diamonds) versus relative forward foot speed at landing (forward foot speed/centre of mass speed) for each step recorded on all terrain types (total 1081 steps). Solid lines are regression fits to the model. The intercept and slope of the fitted line for the compliant jointed model are $0.0252 \pm 0.004$ and $0.0203 \pm 0.010$, respectively ($R^2 = 0.57, p < 0.0001$), and the intercept and slope of the fitted line for the stiff jointed model are $0.120 \pm 0.002$ and $0.056 \pm 0.005$, respectively ($R^2 = 0.29, p < 0.0001$)
