## Supplemental Data 6 for "How human runners regulate footsteps on uneven terrain"

**Figure 6—table supplement 1.** Details of the ANCOVAs performed on the linear model (from main text equation (11)) showing the denominator degrees of freedom, F-value and p-value for the fixed terrain factor, and the estimated slopes $\beta_f$ for the fixed forward foot speed effect.

| dependent variable | factor | DenDF | F-value | p-value | $\beta_f$ |
|---|---|---|---|---|---|
| touchdown leg angle | terrain | 193 | 1.48 | 0.23 | – |
| | fwd. foot speed | 38 | 115.83 | $< 0.0001$ | 0.07 [illegible] |
| fore-aft impulse | terrain | 79 | 1.45 | 0.24 | - |
| | fwd. foot speed | 78 | 12.83 | 0.001 | 0.01 [illegible] |
