## Supplemental Data 7 for "How human runners regulate footsteps on uneven terrain"

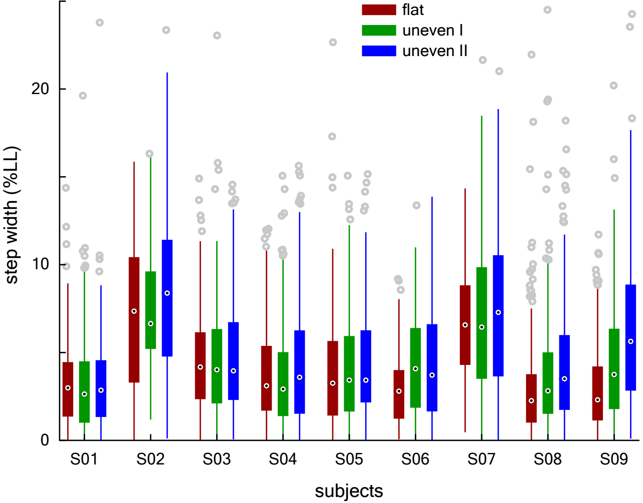


**Figure 7—figure supplement 1. Subject-wise step width statistics.** Step width is expressed as percent leg length (LL). Because the data are skewed with long tails away from zero, we report the median and interquartile range as measures of the central tendency and variability, respectively. Black dots represent medians, boxes show the interquartile range, lines extend to 1.5 times the quartile range, and grey circles represent outliers.
