## Supplemental Data 8 for "How human runners regulate footsteps on uneven terrain"

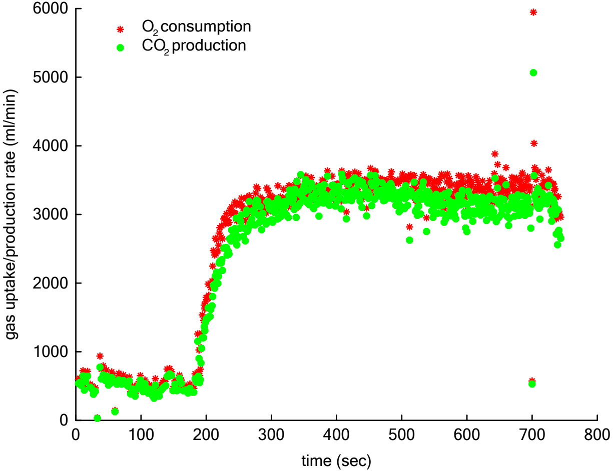


**Figure 7—figure supplement 2. Representative respirometry data.** Breath-by-breath respirometry data for a representative subject running on uneven II. Red stars represent $O_2$ consumption rate and green octagons represent $CO_2$ production rate.
