## Supplemental Data 9 for "How human runners regulate footsteps on uneven terrain"

**Figure 7—table supplement 1.** Kinematic variables on different terrain types reported as mean ± SD, except for meander values which are reported as median ± interquartile range. For each variable, we show details of the ANOVAs performed on the linear model (from main text equation (10)), i.e. the F-value and p-value for the terrain factor. The denominator degrees of freedom for all ANOVAs was 16. Post-hoc comparisons when the ANOVAs reached the significance bound of $\alpha = 0.05$ are reported in the main text results section.

| variable | flat | uneven I | uneven II | F-value | p-value |
|---|---|---|---|---|---|
| net metabolic rate (W/kg) | $13.1 \pm 0.5$ | $13.7 \pm 0.9$ | $13.7 \pm 0.8$ | 2.97 | 0.08 |
| median step width (%LL) | $3.9 \pm 1.9$ | $4.1 \pm 1.5$ | $4.7 \pm 2.0$ | 4.53 | 0.03 |
| IQR step width (% LL) | $3.9 \pm 1.4$ | $4.3 \pm 0.9$ | $5.0 \pm 1.2$ | 3.65 | 0.05 |
| mean step width (%LL) | $4.2 \pm 1.7$ | $4.7 \pm 1.6$ | $5.2 \pm 1.7$ | 8.69 | 0.003 |
| S.D. step width (% LL) | $2.8 \pm 0.8$ | $3.4 \pm 0.6$ | $3.6 \pm 0.6$ | 5.54 | 0.01 |
| mean step length (%LL) | $128 \pm 6$ | $126 \pm 9$ | $125 \pm 9$ | 1.07 | 0.37 |
| S.D. step length (%LL) | $6 \pm 1$ | $7 \pm 4$ | $6 \pm 1$ | 0.64 | 0.54 |
| mean meander ($\times 10^{-4}$) | $3.21 \pm 2.59$ | $3.97 \pm 1.65$ | $4.88 \pm 4.62$ | 1.48 | 0.25 |
| S.D. meander ($\times 10^{-4}$) | $0.67 \pm 0.53$ | $1.33 \pm 1.40$ | $1.27 \pm 2.78$ | 1.58 | 0.23 |
| mean fwd. foot speed (froude num.) | $0.53 \pm 0.17$ | $0.36 \pm 0.10$ | $0.37 \pm 0.12$ | 13.08 | 0.0004 |
| S.D. fwd. foot speed (froude num.) | $0.17 \pm 0.05$ | $0.14 \pm 0.05$ | $0.18 \pm 0.07$ | 1.48 | 0.26 |
| mean CoM speed (m/s) | $3.24 \pm 0.07$ | $3.21 \pm 0.07$ | $3.18 \pm 0.09$ | 2.32 | 0.13 |
| S.D. CoM speed (m/s) | $0.11 \pm 0.03$ | $0.13 \pm 0.04$ | $0.12 \pm 0.03$ | 2.00 | 0.17 |
| mean touchdown leg length (%LL) | $120 \pm 5$ | $119 \pm 4$ | $119 \pm 4$ | 4.28 | 0.03 |
| S.D. touchdown leg length (%LL) | $1.1 \pm 0.7$ | $0.9 \pm 0.3$ | $1.3 \pm 1.2$ | 1.32 | 0.29 |
| mean touchdown leg angle (rad) | $0.20 \pm 0.02$ | $0.20 \pm 0.02$ | $0.21 \pm 0.02$ | 3.90 | 0.04 |
| S.D. touchdown leg angle (rad) | $0.03 \pm 0.02$ | $0.02 \pm 0.003$ | $0.03 \pm 0.02$ | 2.10 | 0.15 |
