## Supplemental Data 10 for "How human runners regulate footsteps on uneven terrain"

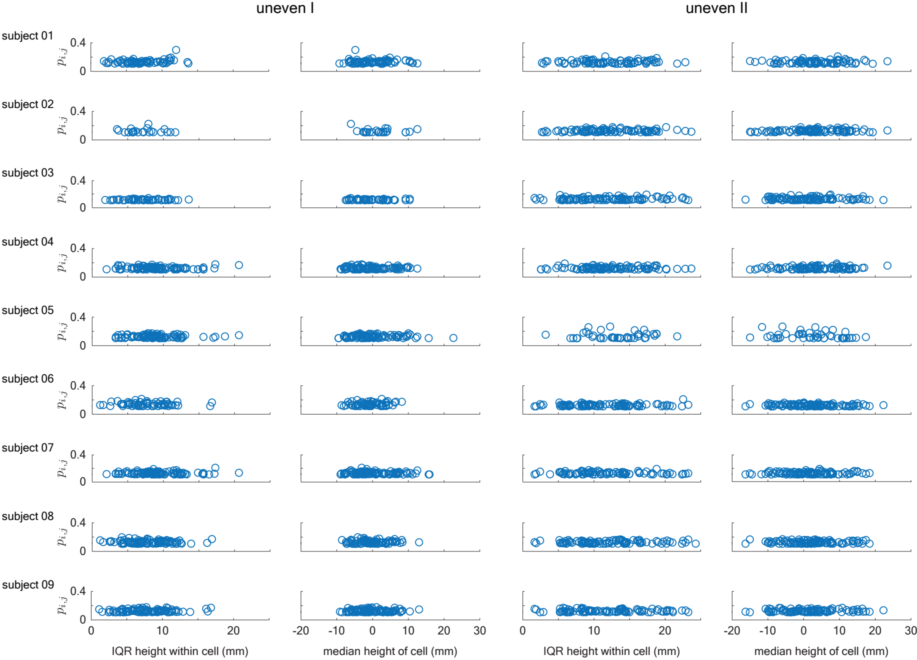


**Table 1—figure supplement 1. Subject-wise foot placement analysis.** Foot placement index $p_{i,j}$ plotted against the median height of the terrain cell and the interquartile range of heights within the terrain cell at landing for all recorded steps on uneven I and uneven II.
